# Osteocyte Differentiation Requires Glycolysis, but Mature Osteocytes Display Metabolic Flexibility

**DOI:** 10.1101/2025.05.09.652291

**Authors:** Matt Prideaux, Mathilde Palmier, Yukiko Kitase, Lynda Faye Bonewald, Tom O’Connell

## Abstract

Recent research has identified metabolic pathways which play key roles in the differentiation and function of osteoblasts and osteoclasts. However, the mechanisms by which osteocytes, the most numerous cells in bone, meet their energetic demands are still unknown. To address this, we used the IDG-SW3 osteocyte cell line to examine changes in metabolism during differentiation from late osteoblasts to mature osteocytes. There was a significant increase in the expression of glycolysis genes (including *Pkm* and *Ldha*), glucose consumption and lactate production during late differentiation of these cells. This was concurrent with the onset of the expression of mature osteocyte markers. Inhibition of glycolysis using the glucose analogue 2-deoxy-d-glucose (2-DG) inhibited IDG-SW3 cell mineralization and differentiation into osteocytes. To examine the effect of glycolysis inhibition on mature osteocytes, we treated differentiated IDG-SW3 cells and long bone osteocytes with 2-DG. Glycolysis inhibition resulted in decreased expression of the bone formation inhibitor *Sost* and mineralization inhibitor *Fgf23*. Concurrently, there was an increase in genes associated with lipolysis (*Lpl*) fatty acid β-oxidation (*Pparδ* and *Cpt1a*). Treatment of differentiated IDG-SW3 cells with the unsaturated fatty acid oleic acid increased *Cpt1a* expression and downregulated *Sost* and *Fgf23*. Application of mechanical stress to IDG-SW3 cells resulted in upregulation of oxidative metabolism, *Pparδ* and *Cpt1a* expression. Long and short chain acylcarnitines were increased in the cortical bone of axially loaded tibiae compared to non-loaded controls, indicative of increased β-oxidation. Overall, our data suggests that while glycolysis is essential for osteocyte differentiation, mature osteocytes are metabolically flexible. Furthermore, β-oxidation may play an important role in the osteocyte response to mechanical stress.

## Introduction

A healthy skeletal system relies upon the coordinated activities of the three main types of bone cells, osteoblasts, osteoclasts and osteocytes. Of these cells, the osteocyte is a master regulator of bone health, by directing the bone forming activity of osteoblasts and the resorptive activity of osteoclasts (^1,2^). However, the location of osteocytes deep within the mineralized matrix has limited their accessibility and hindered research into their biology and function. In particular, while energy metabolism has been shown to have a key role in regulating the function of osteoblasts (^3–6^) and osteoclasts (^5,7,8^), little is known about osteocyte energy metabolism. Given the highly specialized nature of osteocytes, and the challenging conditions of their environment, it would be expected for these cells to have specific requirements in order to meet their energy demands.

Fuel sources for cellular ATP production can come from metabolism of glucose, fatty acids and amino acids and are dependent on nutrient availability, tissue oxygen levels and energy demand (^9^). The process of glycolysis, where glucose is converted into pyruvate via a series of cytoplasmic enzymatic reactions, has been shown to be particularly important for osteoblast (^10–13^) and osteoclast function (^8,14–16^). Under oxygen replete conditions, pyruvate can be further converted into acetyl CoA to fuel the TCA cycle and subsequently oxidative phosphorylation (OxPhos) in mitochondria. Under anerobic conditions, the pyruvate is converted into lactate and released from the cell by monocarboxylate transporters (MCTs) (^17,18^). However, it has been demonstrated that in certain cell types under specific conditions, pyruvate is converted into lactate even in the presence of oxygen. In particular, this is known to occur in cancer cells where it is called the ‘Warburg effect’ (^19^). Recent studies have identified that this process is also important for osteoblast differentiation and mineralization in response to anabolic stimuli such as PTH administration and activation of Wnt/β-catenin signaling (^10,11,20–22^). The role of fatty acids and amino acids in osteoblasts have been less explored, although free fatty acid supplementation has been shown to promote osteoblast differentiation and bone formation during fracture healing (^23^). Additionally, inhibition of fatty acid β-oxidation in osteoblasts *in vivo* resulted in a sustained decrease in bone mass in female mice (^24^).

Despite these recent advances in understanding energy metabolism in osteoblasts, little is currently known about the role of energy metabolism during osteocyte differentiation, or in mature osteocytes. While these cells represent different stages of a common lineage, their functions are highly diverse. Osteoblasts are required to rapidly synthesize and initiate mineralization of the collagenous bone matrix. However, osteocytes act as regulators of bone remodeling in response to mechanical and biochemical signals (^2,25^) and have endocrine functions via signaling to tissues such as kidney and skeletal muscle (^26–28^). Understanding how fuel substrate utilization impacts these processes will provide valuable insights into how long-lived osteocytes are able to survive, particularly in the bone environment in which access to nutrients and oxygen may be restricted compared to other tissues (^29^). In this study we report that glycolysis plays a key role during osteocyte differentiation. However, short-term inhibition of glycolysis in mature osteocytes promotes beneficial effects on regulators of bone remodeling, potentially by upregulating fatty acid β-oxidation. This metabolic switch may play an important role in the response of osteocytes to anabolic stimuli such as mechanical loading.

## Methods

### IDG-SW3 Cell Culture

IDG-SW3 cells were cultured as described in (^30,31^). Briefly, IDG-SW3 cells (passage 12-18) were seeded in collagen-coated 12 or 24 well plates, or T75 culture flasks at a density of 4×10^4^/cm^2^. The cells were cultured at 33°C and 5% CO_2_ in proliferation medium (α-MEM with 10% FBS, 100 U/ml penicillin, 50 µg/ml streptomycin and 50 U/ml IFN-γ) (all Thermo Fisher Scientific, Massachusetts) for 48 hours until fully confluent. The medium was then replaced with differentiation medium (α-MEM containing 10% FBS, 100 U/ml penicillin, 50 µg/ml streptomycin, 50 µg/ml ascorbic acid and 4 mM β-glycerophosphate) and the cells were cultured at 37°C and 8% CO_2_ to promote differentiation. The medium was replaced with fresh differentiation media every 3 days. IDG-SW3 cell experiments were performed with an *n* of 3 or 4 biological replicates per group and each experiment was repeated 2-3 times.

### Inhibition of Glycolysis Using 2-Deoxy-D-Glucose

To determine the effects of glycolysis inhibition on osteocyte differentiation, IDG-SW3 cells were cultured in 12 well plates and differentiated for 9 days as described above. On day 9 the medium was replaced with fresh differentiation medium containing 100 µM - 1 mM 2-deoxy-D-glucose (2-DG, Cayman Chemicals, Michigan) dissolved in PBS. An equivalent volume of PBS was used as the vehicle control. The medium was replaced with fresh differentiation medium containing 2-DG every 3 days until day 20 of differentiation. On day 20 the medium was replaced with fresh differentiation medium, and 2-DG and the cells were harvested 24 hours later (day 21) for analysis.

To determine the effects of glycolysis inhibition on mature osteocytes, IDG-SW3 cells were differentiated for 28 days until they acquired a mature osteocyte phenotype. The cells were then cultured in differentiation medium containing 250 µM 2-DG or PBS vehicle control for 24 hours.

### Quantification of ECM Mineralization

IDG-SW3 cells were differentiated for 21 days in the presence of 100 - 500 µM 2-DG in 24 well plates as described above. On day 21, the cells were fixed in 4% paraformaldehyde for 10 mins at 4°C, rinsed in PBS and stained with 2% Alizarin red (Sigma-Aldrich, Missouri) (pH 4.2) for 5 mins at RT with gentle agitation. The wells were then washed in distilled H_2_O to remove any unbound stain. To quantify the alizarin staining, the dye was extracted from the mineralized matrix in 500 µl 10% cetylpyridinium chloride (Sigma) per well and the optical density was measured at 570 nm (BioTek Synergy HTX, Agilent Technologies, California).

### Cell Death ELISA

To examine whether inhibition of glycolysis affects osteocyte viability, the Cell Death Detection ELISA (Roche, Indiana) was used according to the manufacturer’s protocol, with some adaptations. IDG-SW3 cells were plated in 96 well plates and differentiated for 28 days. On day 28 the medium was replaced with 100 µl of fresh differentiation medium containing PBS or 2-DG (100 µM-1 mM concentration) and incubated for 24 hours. Cells treated with 0.7 mM H_2_O_2_ for 4 hours prior to harvesting were used as a positive control. For harvesting, the medium was removed, and the wells washed 3 times with PBS. 200 µl of incubation buffer was added to each well and incubated for 30 mins at room temperature. The wells were then scraped with a pipette tip, the contents transferred to a microcentrifuge tube and centrifuged. 40 µl of the supernatant was transferred to a fresh tube and diluted 1:5 in incubation buffer. The ELISA procedure was performed as described in the protocol and all steps were performed at room temperature. Briefly, the wells of ELISA plate were coated for 1 hour with 100 µl coating solution and then pre-incubated with 200 µl incubation buffer for 2 hours. The wells were then washed 3 times with 300 µl washing buffer and 100 µl of samples was added to the wells and incubated for 90 minutes. Incubation buffer was used as a blank control. The wells were washed 3 times with washing buffer, 100 µl conjugate solution was added to each well (except the blank controls) and incubated for 90 minutes. The wells were washed 3 times and 100 µl substrate solution was added and incubated for 20 minutes with gentle agitation until color developed. The absorbance was measured at 405 nm. To examine the effects of 2-DG on cell death during osteocyte differentiation, the same procedure was performed on cells cultured with 100 µM-1 mM 2-DG beginning at day 9 of differentiation until day 21.

### Fluid Flow Shear Stress

IDG-SW3 cells were seeded onto type I collagen-coated glass slides (Thermo Fisher) at density of 4×10^4^/cm^2^ and cultured at 33°C and 5% CO_2_ in proliferation medium for 48 hours until confluent. The medium was then replaced with differentiation medium, and the cells were cultured at 37°C and 8% CO_2_ for 28 days for mature osteocyte differentiation. The medium was replaced every 3 days. On day 28, the cells were subjected to steady laminar fluid flow shear stress (FFSS) at a rate of 14 dynes/cm^2^ using the Streamer Gold system (Flexcell International, North Carolina). After 2 hours of FFSS, the cells were either harvested immediately or cultured for a further 3 hours in differentiation medium under static conditions. RNA was extracted by scraping the cells in TriZol (Thermo Fisher) and used for subsequent qRT-PCR analysis. Control cells were treated under identical conditions but were not subjected to FFSS.

### Culture of ex-vivo Osteocyte-enriched Bone

Primary osteocyte-enriched bone pieces were isolated as previously described (^32,33^). Briefly, tibiae, femurs and humeri were aseptically dissected from 3-month-old female C57Bl/6 mice (n=5). The soft tissue was removed, and the marrow flushed with PBS using a 27-gauge needle. The bones were dissected into smaller pieces (approx. 2 mm) and digested sequentially with 2 mg/ml collagenase type IA (Sigma) in α-MEM, 5 mM EDTA /0.1% BSA in HBSS and a final digestion in collagenase for 25 mins at 37°C. The bone pieces were then cultured overnight in primary osteocyte medium (α-MEM with 10% FBS, 100 U/ml penicillin, 50 µg/ml streptomycin). The following day the medium was replaced with fresh medium containing 2 mM 2-DG or PBS as a vehicle control. The bone pieces were then cultured for 24 hours. The bone pieces were snap frozen in LN_2_, pulverized and the pulverized bone powder was stored at −80°C in TriZol (Thermo Fisher) ready for RNA extraction.

### Fatty Acid Treatment of IDG-SW3 Cells

IDG-SW3 cells were cultured in 24 well plates as described previously and differentiated for 28 days into mature osteocytes. Oleic acid (OA, Thermo Fisher) was complexed to fatty acid free bovine serum albumin (BSA, Proliant Biologicals, Boone, Iowa) at a molar ratio of 3:1 by shaking overnight at 37°C. The differentiation medium was removed from the cells and replaced with medium (α-MEM with 10% FBS, 100 U/ml penicillin, 50 µg/ml streptomycin) containing either 100 µM OA-BSA or BSA alone and 400 µM L-carnitine (TCI Chemicals, Portland, Oregan). The cells were cultured for 24 hrs and the lysate was harvested for RNA extraction for PCR analysis.

### RT-PCR

Total RNA was extracted from the IDG-SW3 cell culture lysate or pulverized bone in TriZol using the Direct-zol Miniprep kit (Zymo Research, Irvine, CA) according to the manufacturer’s instructions. 200 ng-1 µg RNA was reverse transcribed into cDNA using the High-Capacity cDNA synthesis kit (Thermo Fisher). Real-time PCR was performed on a Step-One Plus PCR cycler (Applied Biosystems, Massachusetts) using 10 ng of template cDNA. Primer sequences and assays are described in Table 1. *Actb* and *Tbp* were used as housekeeping genes for the IDG-SW3 cultures and *ex vivo* bone chips respectively. Relative expression was calculated using the 2^-ΔΔ^Ct method (Livak, 2001).

### RNA Seq

Library Preparation and Sequencing: To determine changes in metabolism-associated transcripts during osteoblast to osteocyte differentiation, IDG-SW3 cells were differentiated for 4, 9, 18 or 28 days as described previously. RNA was isolated using Trizol as described in (^31^). The concentration and quality of total RNA samples was assessed using Agilent 2100 Bioanalyzer. All samples had a RIN (RNA Integrity Number) of nine or higher. 100 ng of RNA per sample were used to prepare dual-indexed strand-specific cDNA library using KAPA mRNA Hyperprep Kit (Roche). The resulting libraries were assessed for quantity and size distribution using Qubit and Agilent 2100 Bioanalyzer. 200 pM pooled libraries were utilized per flowcell for clustering amplification on cBot using HiSeq 3000/4000 PE Cluster Kit and sequenced with 2×75bp paired-end configuration on HiSeq4000 (Illumina, California) using HiSeq 3000/4000 PE SBS Kit. A Phred quality score (Q score) was used to measure the quality of sequencing. More than 90% of the sequencing reads reached Q30 (99.9% base call accuracy).

Sequence Alignment and Gene Counts: The sequencing data were first assessed using FastQC (Babraham Bioinformatics, Cambridge, UK) for quality control. Then all sequenced libraries were mapped to the mm10 mouse genome using STAR RNA-seq aligner (^34^) with the following parameter: “--outSAMmapqUnique 60”. The reads distribution across the genome was assessed using bamutils (from ngsutils) (^35^). Uniquely mapped sequencing reads were assigned to mm10 refSeq genes using featureCounts (from subread) (^36^) with the following parameters: “-s 2 -p –Q 10”. Quality control of sequencing and mapping results was summarized using MultiQC (^37^). Genes with read count per million (CPM) > 0.5 in more than 3 of the samples were kept. The data was normalized using TMM (trimmed mean of M values) method. Differential expression analysis was performed using edgeR(^38,39^). False discovery rate (FDR) was computed from p-values using the Benjamini-Hochberg procedure.

### IDG-SW3 Seahorse Bioenergetic Assays

IDG-SW3 cells were cultured in a Seahorse XFe96 cell culture microplate (96 wells, Agilent). To ensure their adherence, a collagen coating was performed as follows: 50 µL of collagen solution were added in each well, after 1 h, the solution was removed, and the wells were dried at room temperature for 1 h. This coating step was performed twice. After coating, the cells were plated at a density of 80 000 cells per cm^2^ and incubated at 33 °C in proliferation medium. After 24 h, the medium was replaced with differentiation medium, and the cells were incubated at 37 °C. The medium was changed every 2 to 3 days. IDG-SW3 cell differentiation was delayed in the Seahorse microplates compared with standard culture plates, so the cells were differentiated for 32 days, when the cells were highly mineralized and expressed Dmp1-GFP.

On the day of the experiment, the assay medium - containing Seahorse XF DMEM, 1 mM of Seahorse XF Pyruvate, 5mM of Seahorse XF Glucose and 2mM of Seahorse XF L-Glutamine (Agilent) - was prepared. The inhibitors Oligomycin and Rotenone/Antimycin A were resuspended as recommended by the manufacturer to obtain final concentrations of 1.5 µM and 0.5 µM respectively. The steps of the assay were performed as follows: 1) loading the sensor cartridge with the inhibitors and calibration, 2) rinsing the cells and incubating in the assay medium, 3) measuring ECAR (Extracellular Acidification Rate) and OCR (Oxygen Consumption Rate). Four to six wells per condition were used. The results were analyzed using the Agilent Wave Desktop Software. Immediately after the experiments, the plates containing the cells were frozen at −80°C. Subsequently, the protein content was measured in each well using a BCA assay (Thermo-Fisher) as per the manufacturer’s instructions and the ATP production rate was normalized to the protein content. The results were analyzed using the Agilent Wave Desktop Software.

### NMR Data Collection

To assess changes in metabolites during IDG-SW3 cell differentiation, cells were cultured in T75 flasks and differentiated for up to 28 days. The culture medium and cell lysate were harvested at days 4, 9, 18 and 28 and the medium was changed 24 hours prior to harvest. Metabolites were extracted from the cell layer using methanol/chloroform/water extraction as described in (^31^). For analysis of metabolites in the culture medium, 350 µl of culture medium was run through a 10 KDa MWCO filter (Sigma) to remove protein contaminants. 300 µl of the filtered medium was added to 240 µl deuterated sodium buffer and 60 µl Chenomx reference solution containing 5.0 mM 2,2-dimethyl-2-silapentate-5-sulfonate-d6 sodium salt (DSS-d6) in D_2_O. NMR data were acquired on an Avance III 700MHz NMR spectrometer (Bruker, Billerica, MA) with a TXI triple resonance probe operating at 25C. Spectra were collected with a 1D NOESY pulse sequence covering 12 ppm. The spectra were digitized with 32768 points during a 3.9 second acquisition time. The mixing time was set to 100 ms and the relaxation delay between scans was set to 2.0 seconds.

### NMR Data Processing

The data were processed using Advanced Chemistry Development Spectrus Processor (version 2016.1, Toronto, Canada). The spectra were zero filled to 65536 points, apodized using a 0.3Hz decaying exponential function and fast Fourier transformed. Automated phase correction and 3rd order polynomial baseline correction was applied to all samples. Metabolite concentrations were quantified using the Chenomx NMR Suite (version 8.2, Edmonton, Canada). The DSS-d6 was used as a chemical shift and quantification reference for all spectra and was set to a chemical shift of 0.00 ppm and a concentration of 500 uM. Quantitative fitting of each spectrum was carried out in batch mode, followed by manual adjustment for some spectra to correct for errors arising from spectral overlap.

### Axial Tibial Loading of Murine Bone

7-month-old female C57Bl/6 mice (n=10) were anesthetized using 2.5% isoflurane and their right tibiae were subjected to axial compression under continuous anesthesia for 216 cycles using a sinusoidal waveform of 2Hz (EnduraTEC ELF 3200; Bose, Massachusetts). A peak force of 11N was selected to generate a strain of approximately +1200 με at the medial midshaft of the tibia in C57Bl/6 female 26-week-old mice as described in (^40^). Loading was performed daily for a total of 5 days and after each loading session the mice were returned to their cages, where they were observed to resume unrestricted activity. The left tibia served as a contralateral, non-loaded control. 6 hours after the final bout of loading, the mice were sacrificed, and the tibiae were dissected. Soft tissues and epiphyses were removed, the periosteum was scraped with a scalpel, and the marrow was flushed with PBS using a syringe and 27-gauge needle. The tibiae were then snap frozen in liquid nitrogen and stored at −80°C. To ensure sufficient tissue for metabolic profiling, the tibiae from two mice were combined to provide one sample (n=5 samples per group).

### Metabolic Profiling of Mouse Bone by LC-MS

To extract metabolites from the tissue, the tibiae were homogenized in 500 μl methanol:water (8:2) using a refrigerated microtube homogenizer (BeadBlaster 24R; Benchmark Scientific, New Jersey). The tibiae were weighed prior to homogenization to enable normalization of metabolites to tissue weight. The homogenization tubes were then centrifuged at 10,000 x g for 5 mins at 4°C. 400 μl of supernatant was removed from each homogenization tube and transferred to a 1.5 ml centrifuge tube. A second extraction of each sample was performed by adding 500 μl of methanol:water (8:2) to each homogenization tube and repeating the homogenization procedure. The supernatant from the second extraction was combined with the first extraction and the samples were the evaporated using a SpeedVac SPD1030 vacuum concentrator (Thermo Scientific) at 10 mTorr and 65°C. The dried samples were reconstituted in 25 μl methanol:water (8:2) and 10 μl of sample was used for LC-MS metabolic profiling using the Biocrates MxP Quant 500 kit (Biocrates Inc., Innsbruck, Austria). Data was collected on a Sciex 5500 QTrap (AB Sciex LLC, Framingham, MA)

### Statistical Analysis

Data analysis was performed using GraphPad Prism 9 for Windows (GraphPad Software, La Jolla California USA). Normal distribution was assessed using the Shapiro-Wilk test. Normally distributed data was analyzed by either a two-tailed t-test or One-Way ANOVA with Tukey’s post-hoc test. Non-normally distributed data were assessed using Kruskal-Wallis test with Dunn’s post-hoc test. A p value less than 0.05 was considered significant.

## Results

### Metabolic Profiling During Osteoblast to Osteocyte Differentiation

To examine metabolism changes during osteocyte differentiation we utilized the IDG-SW3 cell line, which differentiates from an osteoblast to mature osteocyte-like phenotype over 28 days in culture (^30,32^). Day 4 (osteoblast), day 9 (early osteocyte), day 18 (mineralizing osteocyte) and day 28 (mature osteocyte) time points were selected for metabolic profiling by nuclear magnetic resonance (NMR) spectroscopy, as these have been shown to represent distinct phenotypic stages (^30,31^). There were few changes in the detected metabolites in the culture medium between day 4 and day 9 of differentiation, although the TCA cycle intermediates citrate and fumarate were elevated at day 9 (Figure 1A, Supplemental Figure 1A and Supplemental File 1)and the branched chain amino acid metabolites methyl-2-oxovalerate, α-ketoisoisovaleric acid and oxo-isocaproate were decreased (Figure 1A and 1B and Supplemental File 1). A significant shift in metabolites was observed on day 18, when the cells are differentiating into osteocytes. There was a dramatic reduction in the medium levels of amino acids including the branched chain amino acids valine, leucine and isoleucine (Figure 1A and 1C) as well as the non-essential amino acids arginine, asparagine, glycine, tyrosine and proline (Figure 1A and Supplemental Figure 1B). Citrate and fumarate levels were strongly reduced compared to day 9, as was succinate (Figure 1A and 1B), suggesting decreased TCA cycle activity. Whereas the increased lactate and decreased medium glucose and pyruvate levels (Figure 1A and 1D) indicate enhanced glycolysis. These changes in amino acids, TCA cycle and glycolysis metabolites were maintained at day 28, although the levels of lactate were lower at day 28 than at day 18. Fewer metabolites were detected in the cell lysates from the IDG-SW3 cells at the same time points (Supplemental Figure 2 and Supplemental File 1). The levels of glutamate, taurine and creatine were increased at days 18 and 28 of differentiation, whereas glycine, glutathione, serine, and creatine phosphate were all decreased in the day 28 differentiated cells.

**Figure 1.**
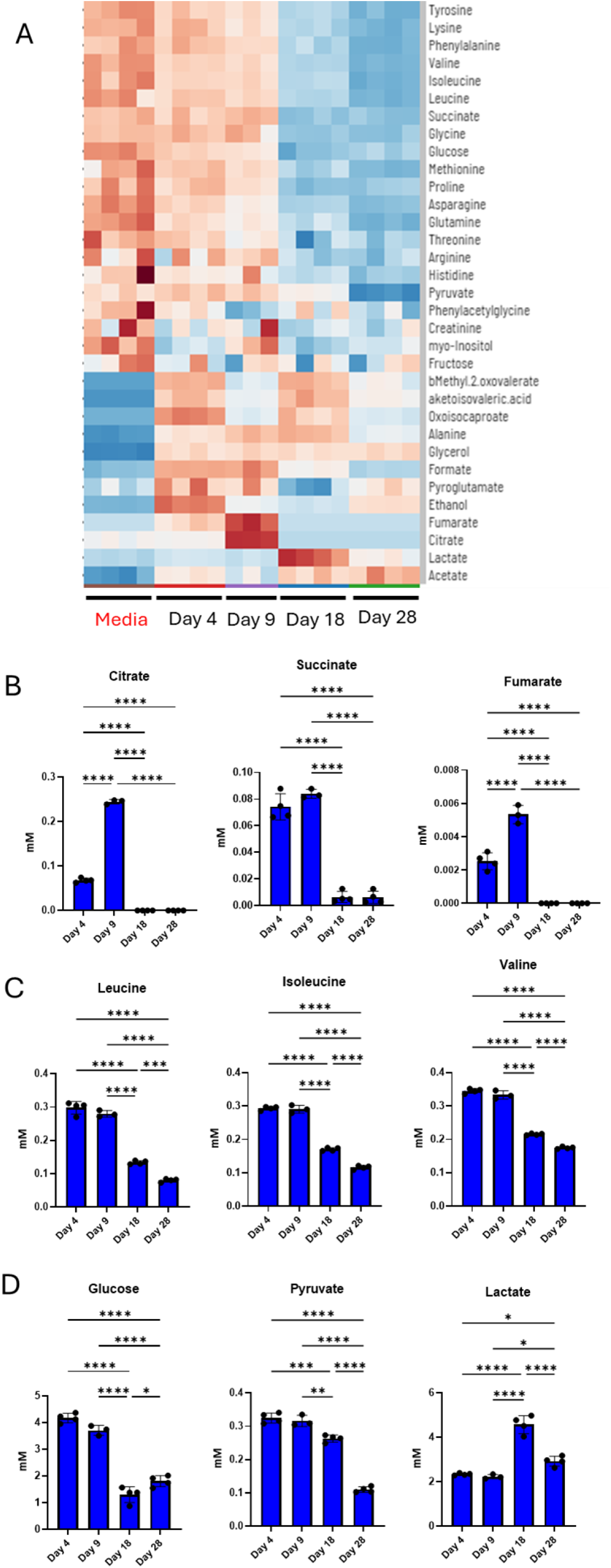
Metabolic profile of conditioned media from IDG-SW3 cells during osteoblast to osteocyte differentiation. (A) Heatmap of metabolites in the media of IDG-SW3 cells at the late osteoblast (day 4), early osteocyte (day 9), mineralizing osteocyte (day 18) and mature osteocyte (day 28) differentiation stage. The Media control was culture media that had not been exposed to cells. (B) Quantification of culture media TCA cycle intermediate metabolites. (C) Quantification of culture media branched chain amino acids. (D) Quantification of culture media glycolysis metabolites (n=3-4 ± SD, ***=p<0.001, **=p<0.01, *=P<0.05. One way ANOVA with Tukey’s post-hoc test).

### Glycolysis is Increased During Osteocyte Differentiation

To further investigate the changes in glycolysis during osteocyte differentiation, we analyzed medium glucose and lactate levels every 3 days during a 27-day IDG-SW3 cell culture time course. There was a significant decrease in medium glucose at day 15 of differentiation compared to day 3 and this was further decreased at days 18-24, before rebounding slightly at day 27 (Figure 2A and Supplemental File 2). The increase in glucose utilization by the IDG-SW3 cells at day 15 was mirrored by increased lactate secretion, with further increases in medium lactate at days 18-24. Interestingly, lactate levels were lower at days 6-12 of differentiation than at day 3, suggesting an increased reliance on oxidative metabolism at this stage of differentiation. To compare the increased glycolysis with osteocyte differentiation, we analyzed the expression of *Dmp1*, *Sost* and *Mepe* and found that the onset of expression of the mature osteocyte markers *Sost* and *Mepe* correlates with that of the increased glycolysis (Figure 2B). *Dmp1* expression, which is an earlier marker of osteocyte differentiation and mineralization (^41^), also peaked concurrently with the increased glycolysis. Furthermore, the expression of the key glycolysis enzyme genes *Hk2*, *Pfkfb3*, *Pkm*, *Pdk1*, *Ldha* and *Slc16a3* showed a similar pattern to that of lactate release and the osteocyte marker genes (Figure 2C).

**Figure 2.**
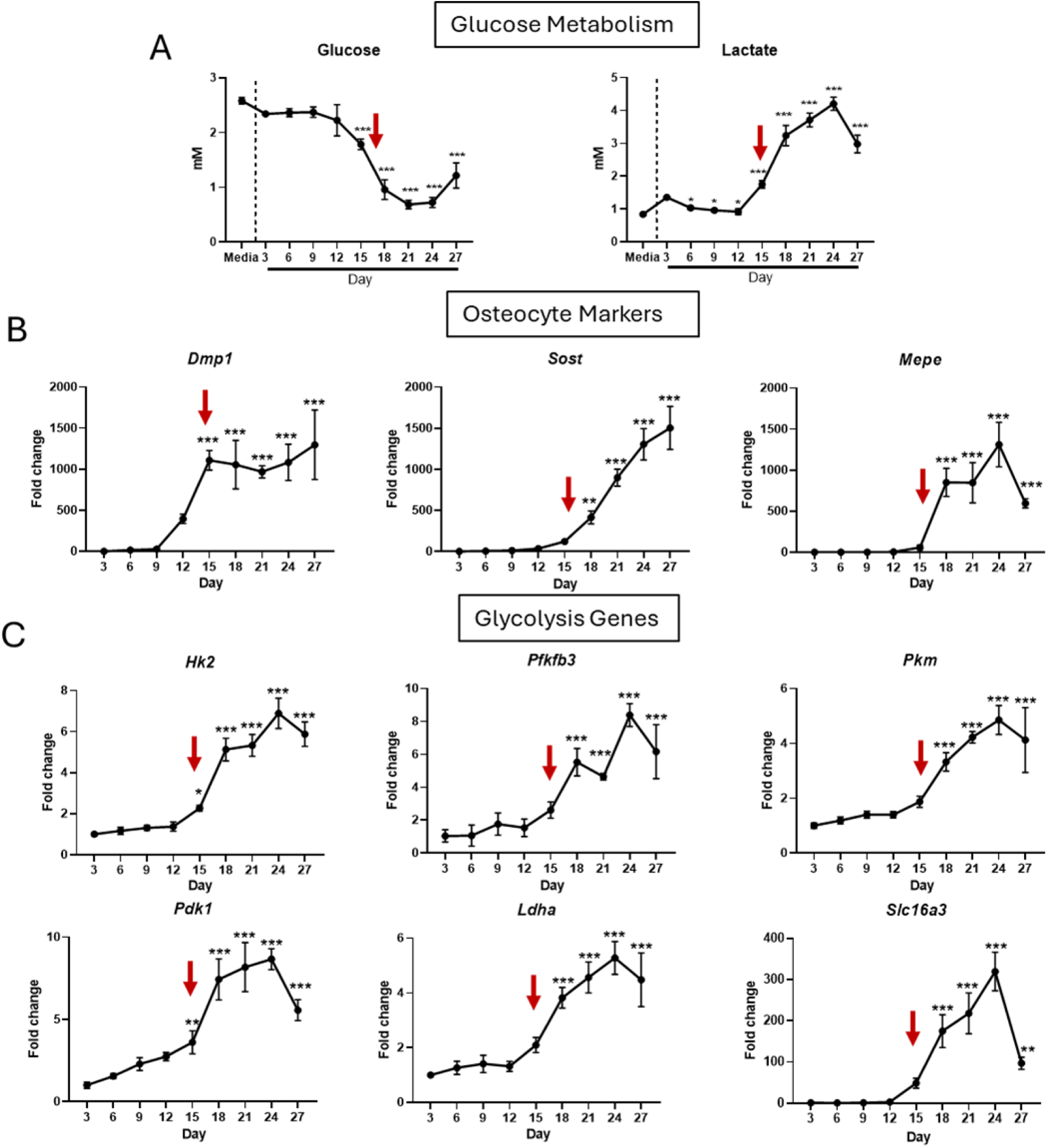
The onset of mature osteocyte marker gene expression correlates with increased glycolysis and expression of glycolysis genes. (A) Media glucose is significantly decreased at day 15 of IDG-SW3 cell differentiation and lactate secretion is significantly increased at the same stage. (B) *Dmp1*, *Sost* and *Mepe* mRNA expression correlates with that of glucose uptake and lactate secretion. (C) The expression pattern of key glycolysis genes mimic those of osteocyte markers and glycolysis (n=3 ± SD, ***=p<0.001, **=p<0.01, *=p<0.05 compared to day 3. One way ANOVA with Tukey’s post-hoc test).

The potent increase in lactate and decrease in glucose led us to further investigate the potential role of glycolysis during osteocyte differentiation. We performed RNA Seq on IDG-SW3 cells at corresponding time points to the metabolite profiling and found a striking increase in the expression of genes involved in glucose metabolism between day 9 and day 18 of differentiation (Figure 3A and B and Supplemental File 3). These included genes involved in glucose uptake (*Slc2a10*), as well as early (e.g., *Hk2*, *Pfkl* and *Pfkp*) and intermediate (e.g., *Gapdh*, *Pgk1* and *Pgam1/2*) steps of the glycolytic pathway. *Pkm,* encoding the enzyme responsible for the generation of pyruvate, was also increased. Furthermore, the elevation of *Pdk1*, which inhibits entry of pyruvate into the TCA cycle combined with the increased expression of *Ldha*, indicates a shift towards pyruvate conversion into lactate instead of oxidation. This is supported by the increased expression of *Slc16a3*, which encodes monocarboxylate transporter 4 (MCT4), an exporter of lactate (^17,18^).

**Figure 3.**
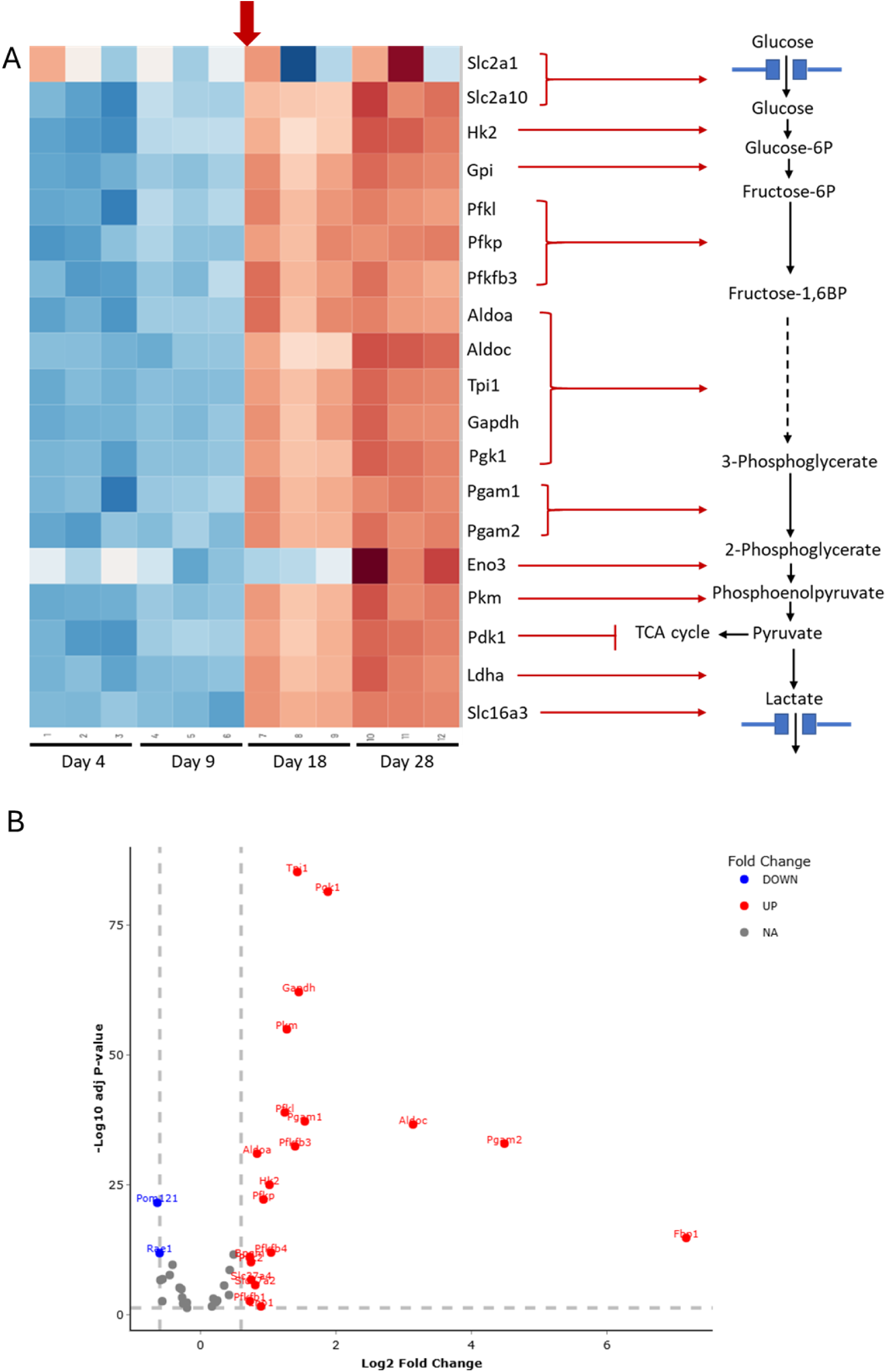
The expression of key glycolysis genes are upregulated during IDG-SW3 cell differentiation. (A) Heatmap of glycolysis genes taken from RNA Seq of differentiating IDG-SW3 cells. (B) volcano plot showing significantly regulated glycolysis genes at day 18 compared to day 9 of differentiation.

### Inhibition of Glycolysis Impairs Osteocyte Differentiation

To determine whether the increased glycolysis plays a functional role during osteocyte differentiation, we treated IDG-SW3 cells with the glucose mimetic 2-deoxy-D-glucose (2-DG), which inhibits the function of hexokinase and glucose-6-phosphate isomerase (^42^) (Figure 4A), starting at day 9 of differentiation. We observed a significant decrease in the expression of the glycolysis enzyme genes *Hk2*, *Pfkfb3*, *Pdk1*, *Ldha* and *Slc16a3* in response to 250 µM 2-DG (Figure 4B). We tested concentrations of 2-DG up to 1 mM and found no effect on cell viability (Figure 3C), however extracellular matrix mineralization was significantly impaired by 2-DG in a dose-dependent manner after 21 days of differentiation (Figure 4D). Consistent with the decreased mineralization, *Alpl*, *Ocn* and *Phex* expression was reduced in cultures treated with 250 µM 2-DG (Figure 4E). 2-DG had no significant effects on the early osteocyte markers *Pdpn* and *Dmp1*, but the mature osteocyte markers *Sost* and *Mepe* were strongly downregulated by 2-DG (Figure 4E). Interestingly, the mRNA expression of the phosphate regulator *Fgf23* was potently elevated by 2-DG treatment (Figure 4E). *Rankl* expression was not affected by 2-DG, however there was a small, but significant decrease in the expression of *Opg*.

**Figure 4.**
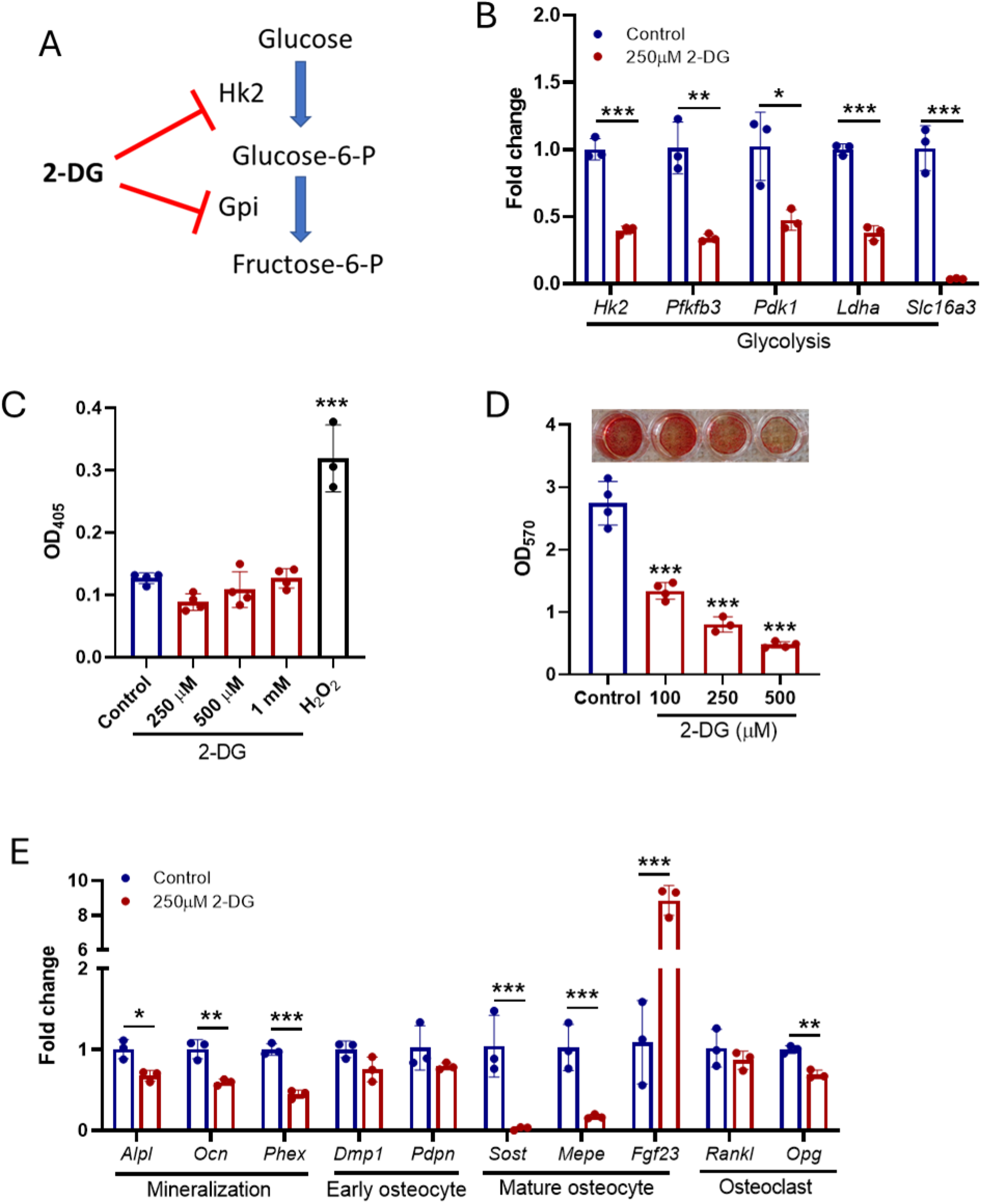
Inhibition of glycolysis during IDG-SW3 cell differentiation inhibits mineralization and osteocyte maturation. (A) 2-deoxy-D-glucose (2-DG) inhibits glycolysis through acting on hexokinase (Hk2) and glucose phosphate isomerase (Gpi). (B) The expression of glycolysis enzyme genes are decreased by 2-DG treatment (C) 2-DG does not induce cell death in the differentiating IDG-SW3 cells. H2O2 was used as a positive control. (D) Extracellular matrix mineralization is dose-dependently impaired by 2-DG. (E) 2-DG decreases the expression of mineralization regulators and mature osteocyte markers but does not affect the expression of early osteocyte markers (n=3-4 ± SD, ***=p<0.001, **=p<0.01, *=p<0.05 compared to day 3. One way ANOVA with Tukey’s post-hoc test or two-tailed T test).

### Glycolysis Regulates Osteocyte Function in Mature Osteocytes

To determine the role of glycolysis in mature differentiated osteocytes, we treated day 28 IDG-SW3 cells with 2-DG for 24 hours and examined the effects on cell viability, osteocyte-expressed regulators of bone remodeling as well as markers of glycolysis and lipid metabolism. Similar to the effects of glycolysis inhibition during osteocyte differentiation, there was no effect of 2-DG on cell viability in the mature osteocytes, whereas the positive control, 0.7 mM H_2_O_2_ induced significant cell death (figure 5A). After 24 hours of treatment with 250 µM 2-DG there was significantly reduced expression of *Sost* and also *Fgf23* (Figure 5B). *Rankl* was increased and *Opg* was decreased, resulting in a significantly elevated *Rankl*/*Opg* ratio in the 2-DG-treated cultures (Figure 5B). Significant decreases were also observed in glycolysis genes (Figure 5C), similar to what we observed with 2-DG treatment during osteocyte differentiation. As 2-DG inhibits the early stages of glycolysis, and therefore the production of pyruvate as fuel for both aerobic glycolysis and the TCA cycle, we hypothesized that the IDG-cells would potentially use fatty acid oxidation as an alternative fuel source. We therefore examined the expression level of a panel of genes involved in lipid synthesis, storage and oxidation. We found increased lipoprotein lipase *(Lpl*) expression (Figure 5D), which is involved in the degradation of circulating triglycerides in lipoproteins (^43^). Fatty acid synthase (*Fasn*), which catalyzes the de novo synthesis of fatty acids (^44^), was decreased by 2-DG treatment. Expression of the rate limiting fatty acid oxidation gene carnitine palmitoyltransferase 1 (*Cpt1a*) was potently increased by glycolysis inhibition, suggesting increased β-oxidation. Additionally, peroxisome proliferator activated receptor delta (*Pparδ*), which is known to activate β-oxidation (^45,46^) was increased whereas *Pparƴ*, which is a promoter of fatty acid synthesis and storage (^47,48^) was unchanged (Figure 5D).

**Figure 5.**
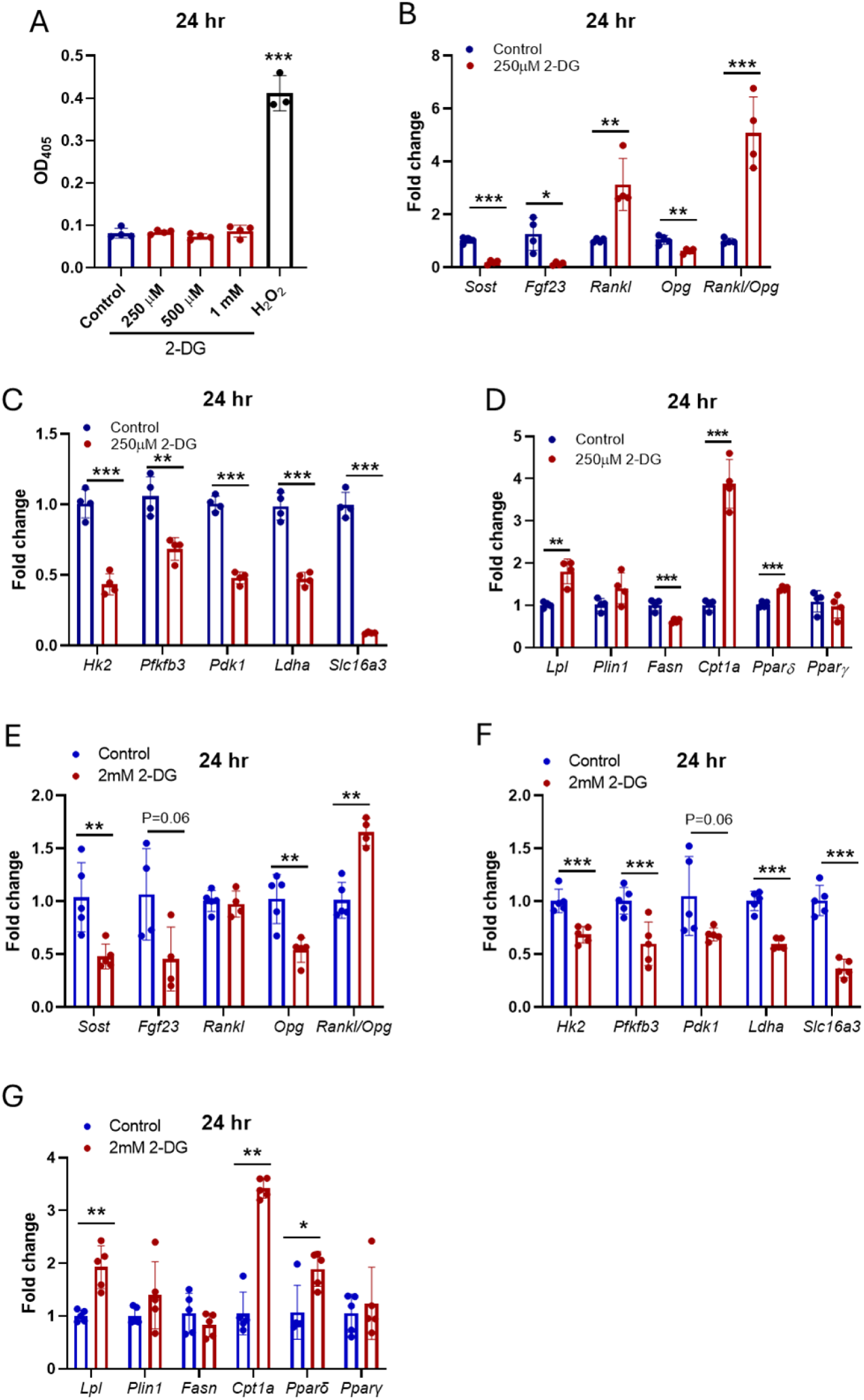
Glycolysis inhibition in mature osteocytes regulates key genes for osteocyte function. (A) 24 hours of 2-DG treatment does not induce cell death in mature IDG-SW3 cell cultures. (B) *Sost*, *Fgf23* and *Opg* expression are decreased by 24 hours 2-DG treatment, whereas *Rankl* is increased, leading to an increased *Rankl*/*Opg* ratio. (C) Glycolysis genes are decreased by 24 hours 2-DG treatment in IDG-SW3 cells. (D) 24 hrs 2-DG treatment regulates lipid metabolism genes in IDG-SW3 cells, particularly those involved in β-oxidation. Similar effects on bone remodelling (E), glycolysis (F) and lipid metabolism (G) genes are observed in osteocyte-enriched bone chip cultures treated with 2 mM 2-DG for 24 hours (n=3-5 ± SD, ***=p<0.001, **=p<0.01, *=p<0.05. One way ANOVA with Tukey’s post-hoc test or two-tailed T test).

To confirm the effects of glycolysis in primary osteocytes, we treated osteocyte-enriched bone chips with 2 mM 2-DG for 24 hours. A higher concentration of 2-DG was required for the primary cultures compared to the IDG-SW3 cells to allow for penetration of the 2-DG through the bone chips to reach the osteocytes. We observed similar effects on bone remodeling genes in the primary osteocytes, with significant decreases in *Sost* and also *Opg*, leading to an elevated *Rankl*/*Opg* ratio (Figure 5E). There was a trend for decreased *Fgf23*, although this did not reach significance. We also saw decreases in glycolysis genes (Figure 5F) and increases in *Lpl*, *Cpt1a* and *Pparδ*, similar to those observed in 2-DG-treated IDG-SW3 cells (Figure 5G).

### Fatty Acid Supplementation Promotes Pathways Involved in Bone Formation and Mineralization

To determine the effect of β-oxidation on osteocyte gene transcription, mature IDG-SW3 cells differentiated for 28 days were treated with 100 µM oleic acid (OA) conjugated to BSA or BSA alone for 24 hrs. OA significantly upregulated *Cpt1a* expression and downregulated *Sost* and *Fgf23* expression (Figure 6). *Rankl*, *Opg* and the *Rankl*/*Opg* ratio were not significantly regulated by 100 µM OA treatment.

**Figure 6.**
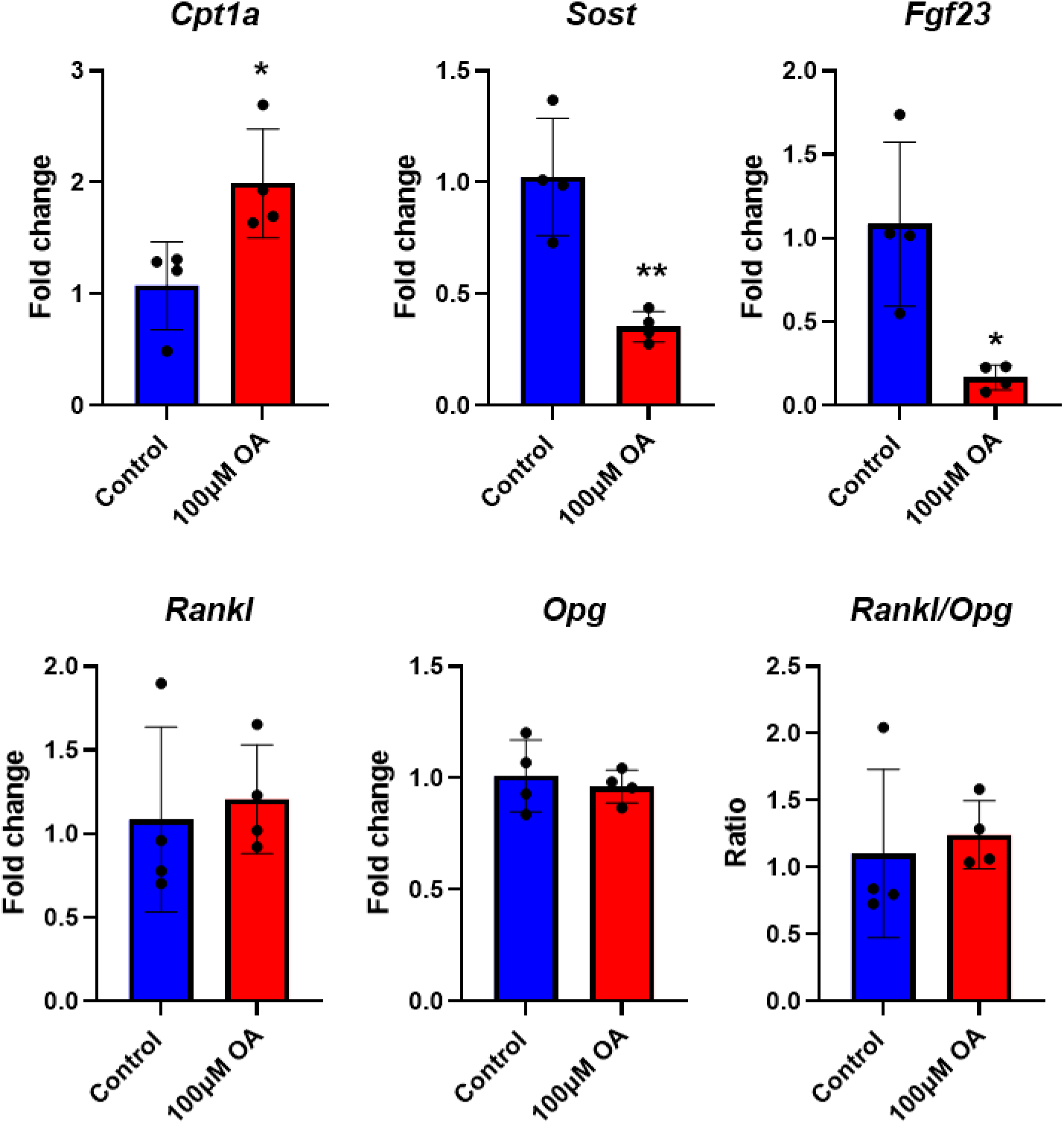
Regulation of bone remodeling genes in mature IDG-SW3 cells by oleic acid. 24 hours of oleic acid (OA) treatment upregulates *Cpt1a* mRNA expression and downregulates *Sost* and *Fgf23*. *Rankl* and *Opg* mRNA expression were unchanged by OA. (n=4 ± SD, **=p<0.01, *=p<0.05 compared to BSA control. Two-tailed T test)

### Mechanical Loading Promotes β-oxidation in Osteocytes

The elevated expression of glycolysis genes in the differentiated IDG-SW3 cells would suggest that glycolysis is a major metabolic pathway for these cells. To confirm this, an ATP rate assay was performed on mature IDG-SW3 cells using the Seahorse system. Surprisingly, almost 90% of the ATP produced by these cells was through oxidative metabolism (Figure 7A). Osteocytes are mechanosensitive cells, and the ATP rate assay protocol involves several washing and mixing steps which would potentially induce fluid flow shear stress (FFSS) on the cells. Therefore, we examined whether FFSS could promote the expression of oxidative metabolism genes and found that expression of the key β-oxidation regulators *Cpt1a* and *Pparδ* were significantly upregulated in mature IDG-SW3 cells subjected to laminar FFSS (14 dynes/cm^2^) (Figure 7B). To examine whether mechanical loading promotes β-oxidation *in vivo*, we performed daily axial loading on the right tibiae of 7-month-old C57Bl/6 female mice for a period of 5 days. Metabolic profiling of the bones showed a distinct separation of metabolites between the loaded and contralateral (non-loaded) tibiae, as can be seen in the principal component analysis (PCA) plot (Figure 7C). Several classes of metabolites were significantly regulated in the loaded bone compared to control, including phosphatidylcholines and lysophosphatidylcholines, ceramides, purines and polyamines, with the majority of these being upregulated in the loaded bone (Supplemental Figure 3 and Supplemental File 4). In particular, we observed a significant increase in the abundance of short chain (SC) and long chain (LC) acylcarnitines, the ratio of SC acylcarnitines to LC acylcarnitines and acetylcarnitine (C2) to free carnitine (C0), markers of β-oxidation (Figure 7D and E).

**Figure 7.**
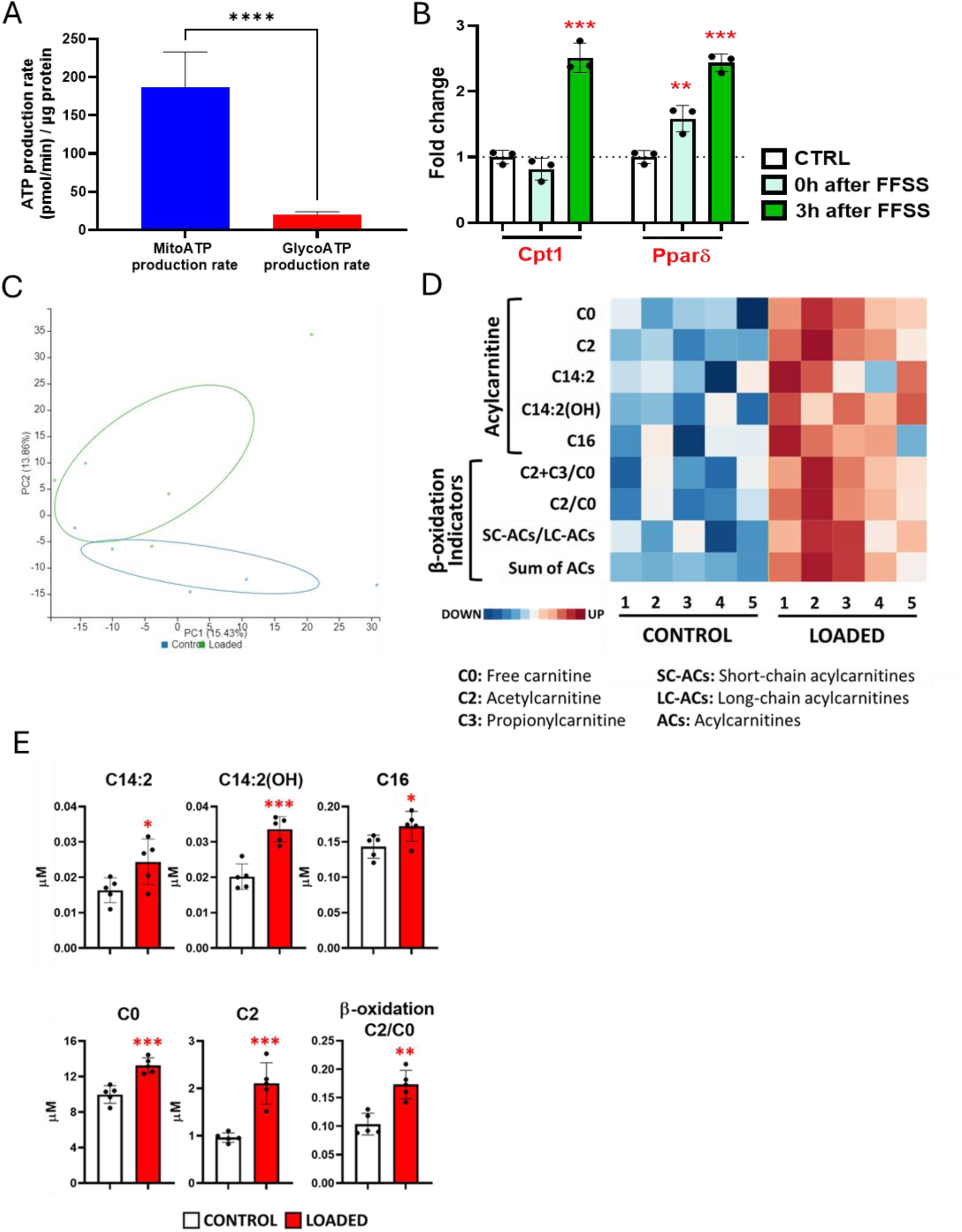
Mechanical loading upregulates markers of β-oxidation in osteocytes and murine cortical bone. (A) ATP production by mature IDG-SW3 osteocytes is primarily oxidative. (B) The expression of *Cpt1a* and *Pparδ* are significantly upregulated following laminar FFSS in matured IDG-SW3 cells. (C) Principal component analysis of metabolic profiling of loaded and control (non-loaded) tibiae. (D) Heatmap showing elevated acylcarnitines and indicators of β-oxidation in loaded compared to control bones. (E) Significantly regulated acylcarnitines in the loaded tibiae (n=3-5 ± SD, ***=p<0.001, **=p<0.01, *=p<0.05. (A) Two-tailed T test. (B) One way ANOVA with Tukey’s post-hoc test).

## Discussion

Despite the importance of osteocytes in regulating bone health, surprisingly little is known about the fuel sources they utilize and how their energy metabolism is regulated during their differentiation. Our studies show that there is a dramatic shift in the metabolic profile of IDG-SW3 cells as these cells differentiate into a mature osteocyte-like phenotype. This is associated with increased utilization of essential and non-essential amino acids as well as increased glucose uptake and lactate secretion. The reason for the increased amino acid usage is unclear, but several of the amino acids are known to play important roles in bone and antagonism or genetic ablation of amino acid transporters has been shown to have detrimental effects on bone formation (^12,49,50^).

We decided to focus on glycolysis due to the striking correlation of the gene expression of glycolytic enzymes and osteocyte markers and the corresponding increase in glucose utilization and lactate secretion. Previous studies have shown that aerobic glycolysis is an essential metabolic pathway for osteoblast differentiation and activity (^10,11,13^) but have not identified the exact cell stage at which this switch to glycolysis occurs. Our data shows that it is the maturing osteocyte that has the highest glycolytic demand, as the glucose uptake and lactate secretion correlate with *Dmp1*, *Sost* and *Mepe* expression. The earlier stages of IDG-SW3 cell differentiation, in which the cells have a late osteoblast-early osteocyte phenotype (^30^), have low glucose utilization and lactate secretion. Furthermore, the levels of the TCA cycle intermediates succinate, citrate and fumarate are higher in the culture media from early differentiated cells, which would suggest greater TCA cycle activity and oxidative metabolism (^51^). However, these metabolites were only detected in the media and not in the cells. This may be due to the sensitivity of NMR to detect these metabolites. Further studies are required to determine the abundance of these intermediates in the early vs late-stage cells.

The importance of glycolysis during osteocyte differentiation was confirmed by inhibition of glycolysis using 2-DG, where we found that decreased glycolysis strongly inhibited the expression of the mature osteocyte markers *Sost* and *Mepe* but had no effect on the early osteocyte marker *Pdpn*. *Fgf23* gene expression was dramatically increased in the 2-DG treated cultures. While FGF23 is well known for its role in regulating serum phosphate via its actions on the kidney (^28^), it has also been shown to directly inhibit mineralization independent of systemic phosphate (^52,53^). Therefore, the elevated *Fgf23* levels may contribute towards the impaired mineralization in the 2-DG treated cultures. Interestingly, although inhibition of glycolysis affected the expression of key bone regulatory genes in the IDG-SW3 cells, it had no effect on cell viability. This suggests that differentiating osteocytes have an inherent metabolic flexibility where they can use alternative energy sources to maintain viability, but at the expense of matrix mineralization and differentiation. The reason why differentiating osteocytes undergo this dramatic increase in glycolysis is yet to be determined, however the embedding of the cells within the mineralized extracellular matrix *in vivo* may be expected to lead to a more hypoxic environment. Hypoxia has been shown to promote glycolysis in tumor cells via the induction of hypoxia-inducible factor 1 (Hif1α) expression (^54^) and this phenomenon also occurs in non-tumor cells (^55^). However, the *in vitro* IDG-SW3 cells were cultured under normoxic conditions, suggesting that there are other factors either instead of, or in addition to hypoxia that regulate glucose metabolism during osteocyte differentiation.

In addition to affecting matrix mineralization and osteocyte differentiation, glycolysis also regulates the function of mature osteocytes *in vitro*. After 24 hrs inhibition of glycolysis in fully differentiated IDG-SW3 cells, *Sost* and *Fgf23* expression was inhibited, which would suggest beneficial effects on bone formation and mineralization. However, the *Rankl*/*Opg* mRNA ratio was also increased after 24 hrs glycolysis inhibition, which would promote osteoclast formation. Similar effects were also observed in primary osteocyte-enriched bone chip cultures. Therefore, inhibition of glycolysis in osteocytes may act to increase bone remodeling through activation of osteoblasts and osteoclasts. This change in the expression of bone regulatory genes was accompanied by a significant downregulation of genes encoding for glycolysis enzymes (e.g., *Hk2*, *Ldha* and *Slc16a3*) and an increase in genes associated with lipid metabolism (*Lpl*, *Cpt1a*, *Pparδ*). This suggests an increase in fatty acid β-oxidation may act as a compensatory mechanism for energy production in response to decreased glycolysis. A similar substrate switching has been observed in acute myeloid leukemia cells in response to glucose deprivation (^56^). The observed increased *Lpl* gene expression suggests a mechanism for the release of fatty acids from triglycerides as a fuel source. PPARδ belongs to the peroxisome proliferator-activated receptor family of transcription factors and has been shown to promote β-oxidation and upregulate the expression of CPT1 in multiple cell types (^57–59^). Deletion of PPARα in osteocytes, another PPAR isoform which promotes β-oxidation, was shown to have a minor effect on bone homeostasis but significantly disrupted whole-body lipid metabolism (^60^).

Treatment of differentiated IDG-SW3 cells with the unsaturated fatty acid oleic acid also resulted in increased expression of *Cpt1a* and downregulation of *Sost*, suggesting that increased β-oxidation may underlie the downregulation of *Sost* in response to glycolysis inhibition. Additional studies are currently ongoing to investigate the role of osteocyte β-oxidation in regulating bone mass.

The regulation of *Fgf23* expression by glycolysis inhibition suggests a novel mechanism by which osteocyte energy metabolism could regulate phosphate homeostasis. The decreased *Fgf23* in response to 2-DG treatment corresponds with an increase in the positive β-oxidation regulators *Pparδ* and *Cpt1a*. Interestingly, it was recently shown that PPARα, negatively regulates the expression of *Fgf23* in UMR106 osteosarcoma cells through the activation of store operated calcium channels (^61^). Lactate has also been shown to promote *Fgf23* expression in these same cells (^62^), therefore decreased aerobic glycolysis (i.e., less lactate production) and increased oxidative metabolism may underlie the downregulation of *Fgf23* in response to glucose inhibition in osteocytes. This is supported by the decreased expression of *Fgf23* in OA treated IDG-SW3 cells.

Surprisingly, when performing bioenergetic assays on the mature IDG-SW3 cells, we observed that these cells were primarily oxidative. We anticipated that these cells would generate most of their energy from glycolysis based on their metabolic and transcriptional profile. A potential explanation for this is that the Seahorse assay protocol requires several media changes and washes. Furthermore, during the assay the plate is mixed several times. All of these steps would lead to mechanical stress on the cells. As one of the main functions of osteocytes is to act as mechanosensors within the bone tissue (^2,26^), this mechanical stress on the IDG-SW3 cells may affect their metabolism. Indeed, when we subjected the cells to laminar FFSS we found that both *Pparδ* and *Cpt1a* were upregulated within 3 hours. Therefore, we propose that while sedentary, unstimulated osteocytes are predominantly glycolytic, mechanically loaded cells switch to a more oxidative state which requires β-oxidation of fatty acids.

While our in vitro experiments demonstrated that osteocytes are metabolically flexible and utilize both glucose and fatty acids, it is impossible to fully replicate the osteocyte environment *in vivo*. To investigate osteocyte metabolism *in vivo*, we performed metabolic profiling on the cortical bone of mechanically loaded or control tibiae from adult female mice. We found significant upregulation of SC and LC acylcarnitines in the loaded tibiae, as well as indicators of increased β-oxidation. It is important to note that this bone does not exclusively contain osteocytes, and there will be a contribution from other cell types in this bone metabolic profile. Fatty acid β-oxidation has been shown to be important for the function of both osteoblasts (^63,64^) and osteoclasts (^65,66^) and our data cannot exclude a role for β-oxidation in the mechanoresponse of these cells. However, the *in vivo* results are consistent with the increased *Cpt1a* in the FFSS treated IDG-SW3 cells and the switch to an oxidative state that we observed during the bioenergetic assays. This increase in oxidative metabolism may be a mechanism to provide additional ATP in order to meet an elevated energy demand. Fatty acid oxidation provides a greater ATP yield than pyruvate oxidation or glycolysis (^67^) and ATP generation and release is an essential component of the osteocyte mechanoresponse (^68–70^). Interestingly, β-oxidation has been shown to decline in many different tissues with aging (^71,72^) and the anabolic response of bone to loading is also impaired with age (^73,74^). Future studies will investigate whether impaired β-oxidation in osteocytes contributes towards the age-associated decline in bone health.

In summary, we have shown that glycolysis is essential for osteocyte differentiation and the acquisition of a mature osteocyte phenotype. However, mature osteocytes display a high degree of metabolic flexibility to maintain viability. While quiescent osteocytes appear to primarily use glycolysis, mechanically stressed cells become more oxidative, with an increased reliance on fatty acid β-oxidation. Therefore, promoting β-oxidation in osteocytes may mimic some of the beneficial effects of mechanical loading on these cells and potentially open a new therapeutic avenue for bone health.

## Acknowledgements

The authors thank The Center for Medical Genomics at Indiana University for performing RNA sequencing analysis and the Metabolomics Initiative at the Indiana Center for Musculoskeletal Health for metabolomics analysis. We would like to acknowledge funding support from NIH R01AG076569 (MPr and YK), P01AG039355 (LFB and TOC). MPa received a scholarship from the Fulbright Program.

## Author Contributions

MPr, LFB and TOC designed the study and concept. MPr, MPa, YK and TOC performed the experiments and analyzed the data. MPr, MPa and TOC wrote the original manuscript. MPr, MPa, YK, LFB and TOC revised and edited the manuscript. All authors read and approved the final manuscript.

## Data Availability

All of the data that support the findings of this study are available from the corresponding author on request. RNA sequencing raw data will be deposited in Gene Expression Omnibus (GEO) and made freely available.

## Supplemental Figure Legends

**Supplemental Figure 1.**
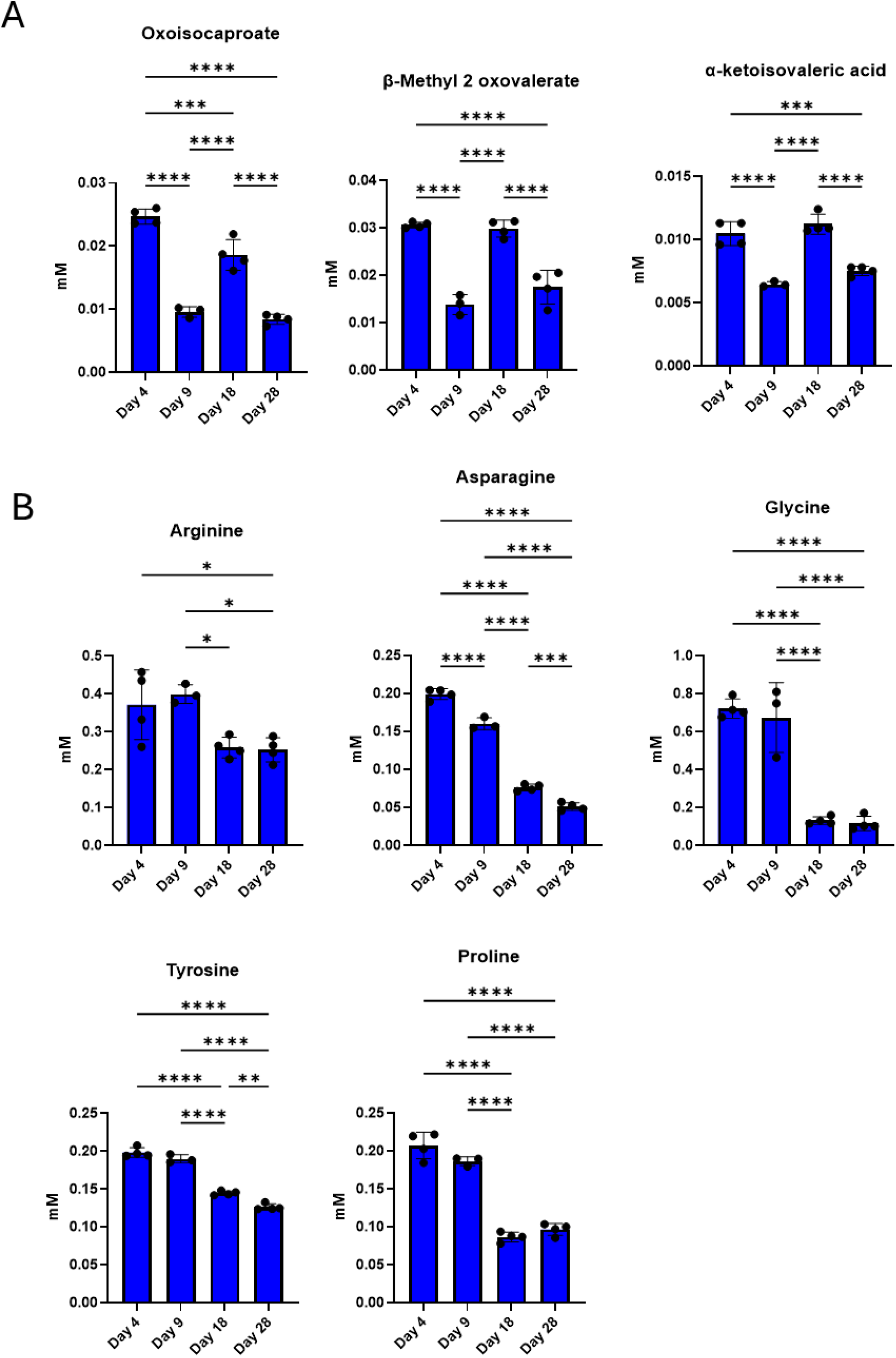
Metabolic profile of cell culture media metabolites during IDG-SW3 cell differentiation. (A) Quantification of culture media branched chain amino acid metabolites. (B) Quantification of culture media non-essential amino acids. (n=3-4 ± SD, ***=p<0.001, **=p<0.01, *=P<0.05. One way ANOVA with Tukey’s post-hoc test).

**Supplemental Figure 2.**
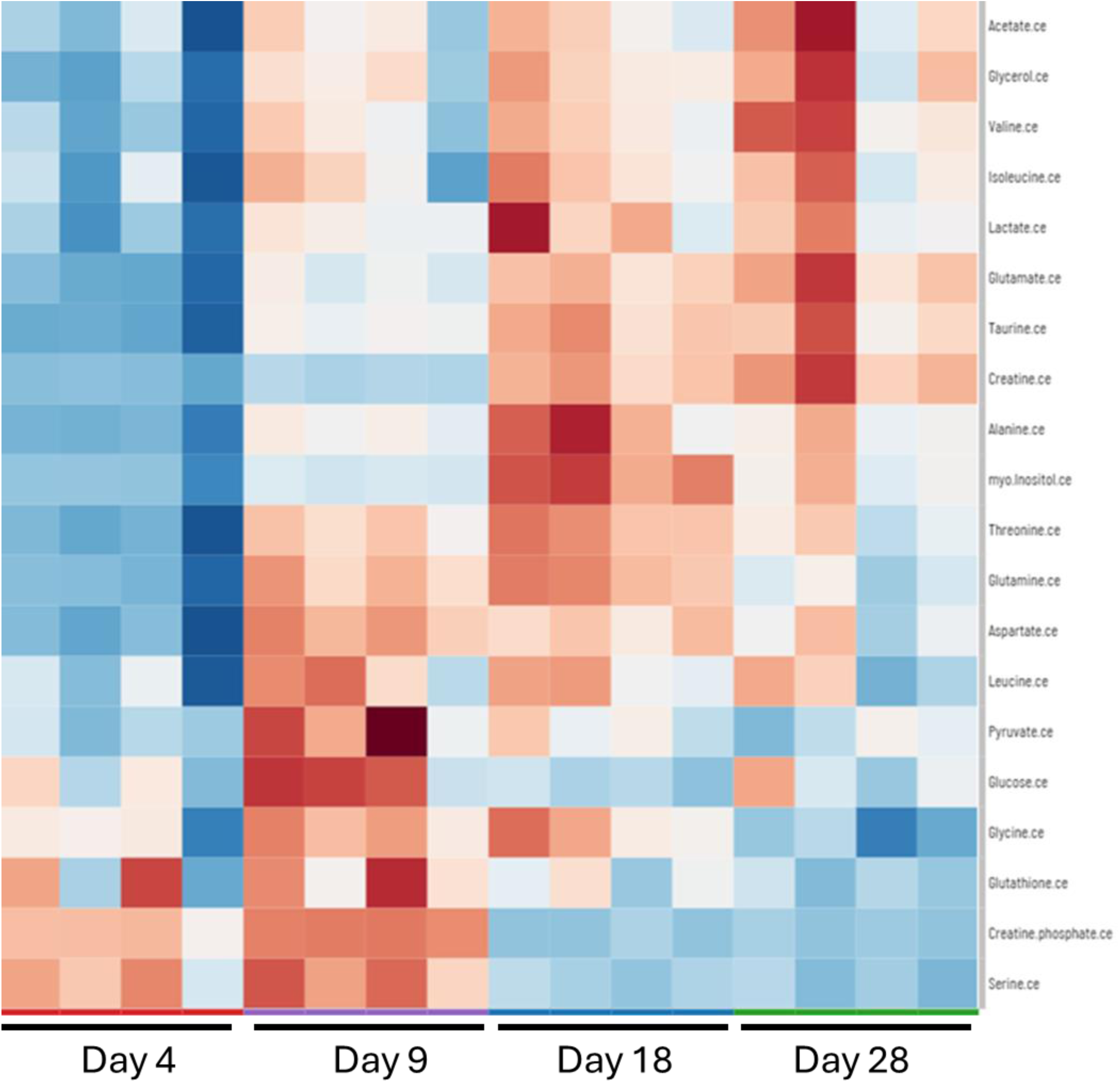
Heatmap of metabolites in IDG_SW3 cell culture lysate that are significantly changed during differentiation into osteocytes.

**Supplemental Figure 3.**
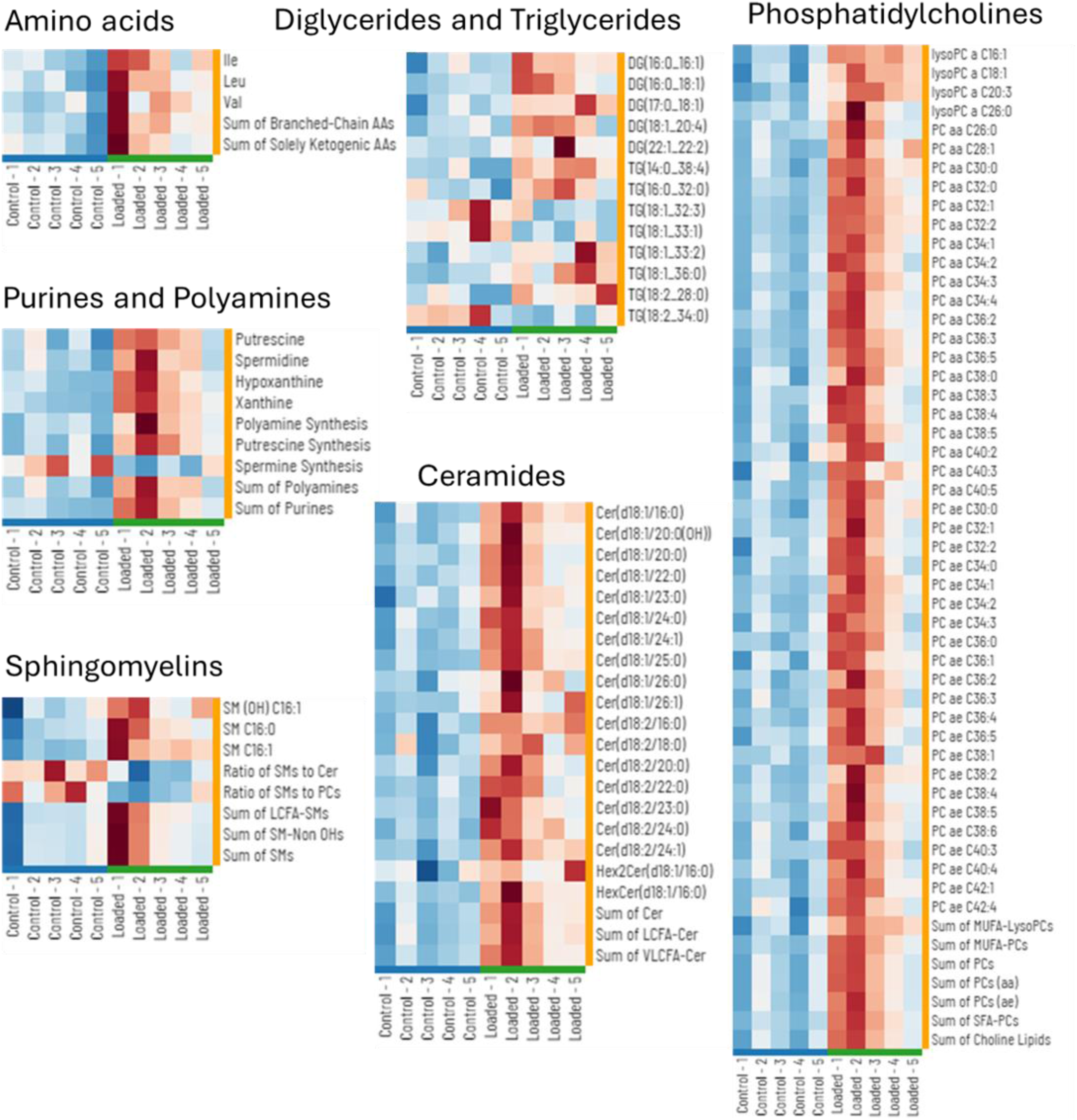
Heatmaps from metabolic profiling of non-loaded (control) and loaded mouse tibiae (additional data from **Figure 7**).

## Supplemental File Information

**Supplemental File 1.** Data from NMR profiling of IDG-SW3 cell culture media during differentiation into osteocytes (from Figure 1 and Supplemental Figure 1).

**Supplemental File 2.** Data from NMR profiling of IDG-SW3 cell culture lysate during differentiation into osteocytes (from Supplemental Figure 2).

**Supplemental File 3.** RNA sequencing data showing differentially expressed genes during IDG-SW3 differentiation (from Figure 3).

**Supplemental File 4.** Biocrates MxP Quant500 metabolic profiling data from non-loaded and loaded mouse tibiae (from Figure 7 and Supplemental Figure 3)

